# In Vivo Emergence of a Novel Protease Inhibitor Resistance Signature in HIV-1 Matrix

**DOI:** 10.1101/865840

**Authors:** Rawlings Datir, Steven Kemp, Kate El Bouzidi, Petra Mlchocova, Richard Goldstein, Judy Breuer, Greg J. Towers, Clare Jolly, Miguel E. Quiñones-Mateu, Patrick S. Dakum, Nicaise Ndembi, Ravindra K. Gupta

## Abstract

**Background:** Protease Inhibitors (PIs) are the second- and last-line therapy for the majority of HIV-infected patients worldwide. Only around 20% of individuals who fail PI regimens develop major resistance mutations in protease. We sought to explore the role of mutations in *gag-protease* genotypic and phenotypic changes within six Nigerian patients who failed PI-based regimens without known drug resistance associated *protease* mutations in order to identify novel determinants of PI resistance.

**Methods:** Target enrichment and NGS by Illumina Miseq were followed by haplotype reconstruction. Full length *gag*-protease regions were amplified from baseline (pre-PI) and virologic failure (VF) samples, sequenced and used to construct *gag*/*protease* pseudotyped viruses. Phylogenetic analysis was performed using maximum likelihood methods. Susceptibility to lopinavir (LPV) and darunavir (DRV) were measured using a single-cycle replication assay. Western blotting was used to analyse Gag cleavage.

**Results:** In one of six participants (subtype CRF02_AG) we found 4-fold lower LPV susceptibility in viral clones during failure of second line treatment. A combination of four mutations (S126del, H127del, T122A and G123E) in p17 matrix of baseline virus generated a similar 4x decrease in susceptibility to LPV but not darunavir. These four amino acid changes were also able to confer LPV resistance to a subtype B gag-protease backbone. Western blotting did not demonstrate significant Gag cleavage differences between sensitive and resistant isolates. Resistant viruses had around 2-fold lower infectivity compared to sensitive clones in the absence of drug. NGS combined with haplotype reconstruction revealed resistant, less fit clones emerged from a minority population at baseline and thereafter persisted alongside sensitive fitter viruses.

**Conclusions:** We have used a multi-pronged genotypic and phenotypic approach to document emergence and temporal dynamics of a novel protease inhibitor resistance signature in p17 matrix, revealing the interplay between Gag associated resistance and fitness.

## Introduction

As antiretroviral therapy (ART) scale up progresses globally in the absence of universal viral load monitoring, significant numbers of persons living with HIV (PLWH) are experiencing virological failure (VF) with emergent drug resistance(1–3). In addition, pre-treatment drug resistance (PDR) has been rising over the past decade(4–6). Although integrase inhibitors are now recommended by WHO in regions where PDR exceeds 10%(7, 8), second line ART in low and middle-income countries (LMIC) is likely to remain dependent on boosted protease inhibitors (PI), specifically lopinavir/ritonavir or atazanavir/ ritonavir.

Studies demonstrate that the detection of major canonical protease mutations is around 20% in PLWH treated with P-containing combination ART(9, 10), raising the question of how virologic failure occurs in the remaining cases. Inadequate adherence to medication has been implicated(11–13), but determinants of susceptibility outside the protease gene have also been considered(14). Interestingly, although PI monotherapy can be effective in some populations in clinical practice (15), this is associated with a higher prevalence of major PI resistance mutations at VF as compared to PI combined with 2 two nucleoside reverse transcriptase inhibitors (NRTI)(16, 17).

The HIV-1 envelope (Env) has been reported in two studies to impact PI susceptibility (18, 19), with a number of reports of diverse *env* sequence changes during PI failure(20, 21). Gag is highly polymorphic across HIV-1 subtypes, and existing literature reports diverse mutations occurring both within and outside cleavage sites following treatment with older PI such as indinavir, saquinavir and nelfinavir in subtype B infections (14, 20–24). Although there has been very limited evidence on the role of HIV-1 *gag* in susceptibility to modern boosted protease inhibitors such as lopinavir/ritonavir used in second line ART in non-B subtypes, we and others have reported that around 1 in 6 individuals infected with non-subtype B HIV who fail modern PI have Gag encoded reduced phenotypic susceptibility to PI(25–29), though specific amino acid determinants have remained elusive.

Cleavage site mutations are thought to partially restore efficient cleavage by protease in the presence of bound drug(30, 31). The mechanism for non-cleavage site mutations may include allosteric changes in protease-gag interactions that influence the efficiency by which protease locates cleavage sites through dynamic intermolecular interactions in the presence of drug(32, 33). For example, our group previously reported the emergence of T81A in Gag that appeared to correlate with reduced susceptibility to the modern PI lopinavir in a subtype AG infected individual in France(26). This mutation was predicted to impact intermolecular interactions between Gag and protease by Deshmukh and colleagues using nuclear magnetic resonance (NMR)(33).

Here we sought to explore the role of mutations in *gag* encoded determinants of reduced PI susceptibility in non-subtype B HIV-1 and to elucidate their evolution in people living with HIV (PLHIV) in Nigeria.

## Methods

### Study participants

We identified six individuals on second line, protease inhibitor based ART who experienced virological failure without major protease mutations from a PEPFAR funded treatment cohort in Nigeria and who had samples collected on at least two time-points (pre second-line initiation and following second-line virologic failure). Having previously reported that baseline phenotypic susceptibility was not associated with subsequent virologic ‘failure’ (34) in this cohort, here we sought to explore changes over time in phenotypic susceptibility that could be associated with changes in HIV-1 *gag*-protease

### Next generation sequencing

Manual nucleic acid extraction was done using the QIAamp Viral RNA mini kit, Qiagen (Hilden, Germany) with a plasma input volume of 0.5-1.5 mL. The first strand of cDNA was synthesised using SuperScript IV reverse transcriptase, Invitrogen, (Waltham, MA, USA), followed by NEBNext second strand cDNA synthesis E6111, New England Biolabs GmbH, (Frankfurt, Germany). Sample libraries were prepared as per the SureSelect^XT^ automated target enrichment protocol, Agilent Technologies (Santa Clara, CA, USA) with in-house HIV baits. Whole genome deep sequencing was performed using Illumina Miseq platform (San Diego, CA, USA). Trimmed reads were then compared to a reference panel of 170 HIV subtypes/CRFs from the Los Alamos database (https://www.hiv.lanl.gov) and the best match was used for reference mapping. Duplicate reads were removed from the BAM files and a consensus sequence was generated using a 50% threshold. Mutations were included if they were present at over 2% frequency within the read mixture at that position, with a minimum read depth of 100. An in-house custom script was used to identify SNPs at each position by BLAST analysis of individual HIV *pol* against the HXB2 reference genome.

### Haplotype Reconstruction and Phylogenetics

Whole-genome haplotype reconstruction was performed using a newly developed maximum-likelihood method, HaROLD (Haplotype assignment of virus NGS data using co-variation of variant frequencies, Richard A. Goldstein, Asif U. Tamuri, Sunando Roy, Judith Breuer bioRxiv 444877; doi: https://doi.org/10.1101/444877). SNPs were assigned to each haplotype so that the frequency of a variant at any time point was represented by the sum of the frequencies of the haplotypes containing that variant. Time-dependent frequencies for longitudinal haplotypes were optimised by maximizing the log likelihood, which was calculated by summing over all possible assignments of variants to haplotypes. Haplotypes were then reconstructed based on posterior probabilities. The calculations were repeated with a range of possible haplotype numbers and the optimal number of haplotypes was determined by the resulting value of the log-likelihood. After constructing haplotypes, a refinement process remapped reads from BAM files to the constructed haplotypes. Haplotypes were also combined or divided according to AIC scores, in order to give the most accurate representation of viral populations. Phylogenetic trees of constructed haplotypes were constructed using RAxML-NG using the GTR model and 1000 bootstraps.

### Coreceptor usage

CCR5 / CXCR4 usage was predicted using *env* sequences with the online tools *Geno2Pheno* (https://www.geno2pheno.org) and WebPSSM (https://indra.mullins.microbiol.washington.edu/webpssm/).

### Amplification of full-length gag-protease genes

We amplified from plasma taken before PI initiation and from a failure time point for each individual. NGS was used to obtain a consensus whole genome sequence for each of the 12 samples. Full length *gag-protease* sequences were obtained from plasma by standard PCR: HIV-1 RNA was extracted from plasma samples using the QIAamp viral RNA extraction kit. Using previously described techniques,(35, 36) full-length gag-protease was amplified and cloned into a subtype B-based (p8.9NSX+) vector. Clonal sequencing of up to 10 plasmids was performed by standard Sanger sequencing. The variant that most closely represented the next-generation sequencing derived consensus, was taken forward for phenotypic testing. Sequences were manually analysed using DNA dynamo software (http://www.bluetractorsoftware.co.uk) software. Protease sequences were analysed for PI resistance mutations using the Stanford Resistance Database (https://hivdb.stanford.edu). Phylogenetic analysis was performed using maximum likelihood methods in MEGA v7.0 (37) Bootstrapping was performed as previously described(26).

### Site directed mutagenesis

Site directed mutagenesis was carried out using Quick Change^TM^ (Stratagene) according to manufacturer instructions. Mutagenesis was verified by Sanger sequencing.

### PI susceptibility and infectivity assays

PI susceptibility and viral infectivity were determined using a previously described single assay. Briefly, 293T cells were co-transfected with a Gag-Pol protein expression vector (p8.9NSX) containing cloned patient-derived full-length gag-protease sequences, pMDG expressing vesicular stomatitis virus envelope glycoprotein (VSV-g), and pCSFLW (expressing the firefly luciferase reporter gene with the HIV-1 packaging signal) as previously described. PI drug susceptibility testing was carried out as previously described(35). Transfected cells were seeded with serial dilutions of lopinavir and harvested pseudovirions were used to infect fresh 293T cells. To determine strain infectivity, virus was produced in the absence of drug.

Infectivity was monitored by measuring luciferase activity 48 h after infection. Results derived from at least two independent experiments (each in duplicate) were analysed. The IC50 was calculated using GraphPad Prism 5 (GraphPad Software Inc., La Jolla, CA, USA). Susceptibility was expressed as a fold change in IC50 as compared to the subtype B reference plasmid p8.9NSX. Replicative capacity of these viruses was assessed by comparing the luciferase activity of recombinant virus with that of the WT subtype B control virus in the absence of drug. Equal amounts of input plasmid DNA were used, and it has previously been shown that percentage infectivity correlates well with infectivity/ng p24 in this system(35). Differences in PI susceptibility were compared with the paired t test. The PI drugs used in this study were obtained from the AIDS Research and Reference Reagent Program, Division of AIDS, NIAID, NIH.

### Western Blot Analysis

Using a previously described method(38), equal amounts of each of the viral clone plasmid was used to transfect 293T cells, in addition to a VSV-G plasmid and reporter genome expressing plasmid. Each of the pseudovirions was produced in the absence and presence of a range of concentrations of LPV, added 16 hours following transfection.

Forty-eight (48) hours post transfection with the plasmid preparations, the culture supernatant was harvested and passed through a 0.45-μm pore-size filter to remove cellular debris. The filtrate was centrifuged at 14,000rpm for 90 minutes to pellet virions. The pelleted virions were lysed in Laemmli reducing buffer (1M Tris-HCl pH 6.8, SDS, 100% glycerol, β-mercaptoethanol and bromophenol Blue). Cell lysates were subjected to electrophoresis on SDS, 4–12% Bis-Tris Protein Gels (Thermo Fisher Scientific) under reducing conditions. This was followed by electroblotting onto PVDF membranes. The HIV-1 Gag proteins were visualized by a trans-Illuminator (Alpha Innotech) using anti-p24 Gag antibody.

### Ethics

Informed consent was obtained from all participants and ethics approval for virological testing was obtained from the Nigeria National Research Ethics Committee of Nigeria (NHREC/01/01/2007). Ethical approval was also obtained from the ethics board of UCL, UK.

## Results

### Phenotypic drug susceptibility following PI failure

Participant characteristics of the six HIV-infected individuals failing PI based second line ART are shown in figure Table 1. Three were infected with CRF02_AG recombinant strain and three with subtype G HIV strains. Phenotypic PI susceptibility testing was performed on plasma derived clones obtained at two time points – before PI treatment (baseline) and at virologic failure (VF).

**Table 1:** Participant and virus characteristics. VF: viral failure

| Patient<br>(Subtype) | Timepoint | Viral<br>Load | Time<br>between<br>Baseline<br>and VF<br>Sample<br>(Months) |
| --- | --- | --- | --- |
| Patient 1<br>(CRF02_AG) | Baseline | 140,991 | 34 |
|  | VF | 6,193 |  |
| Patient 2<br>(G) | Baseline | 20,178 | 42 |
|  | VF | 117,942 |  |
| Patient 3<br>(CRF02_AG) | Baseline | 271,974 | 50 |
|  | VF | 74,224 |  |
| Patient 4<br>(CRF02_AG) | Baseline | 24,693 | 36 |
|  | VF | 32,683 |  |
| Patient 5<br>(G) | Baseline | 274,504 | 31 |
|  | VF | 16,304 |  |
| Patient 6<br>(CRF02_AG) | Baseline | 18,056 | 64 |
|  | VF | 66,277 |  |

Participant 6 had a significant difference in PI susceptibility between baseline and failure time points (Figure 1 and supplementary figure 1). At VF, the LPV IC50, expressed as fold change (FC) compared to the subtype B reference was 20.3 compared to 5.2 prior to initiation of LPV treatment. We phenotyped four clones from baseline, all with similar LPV susceptibility. Baseline genotype (pre PI) indicated that the individual had developed extensive resistance to first line ART with nucleoside reverse transcriptase inhibitor (NRTI) mutations K65R, M184I conferring high level tenofovir and lamivudine resistance respectively, as well as K103N and Y181C conferring to non-nucleoside reverse transcriptase inhibitors (NNRTI). The co-occurrence of the latter two non-NRTI mutations suggests that the individual may have been pre-treated with first line ART containing nevirapine, or have received single dose nevirapine for prevention of mother to child transmission(39).

**Fig 1:**
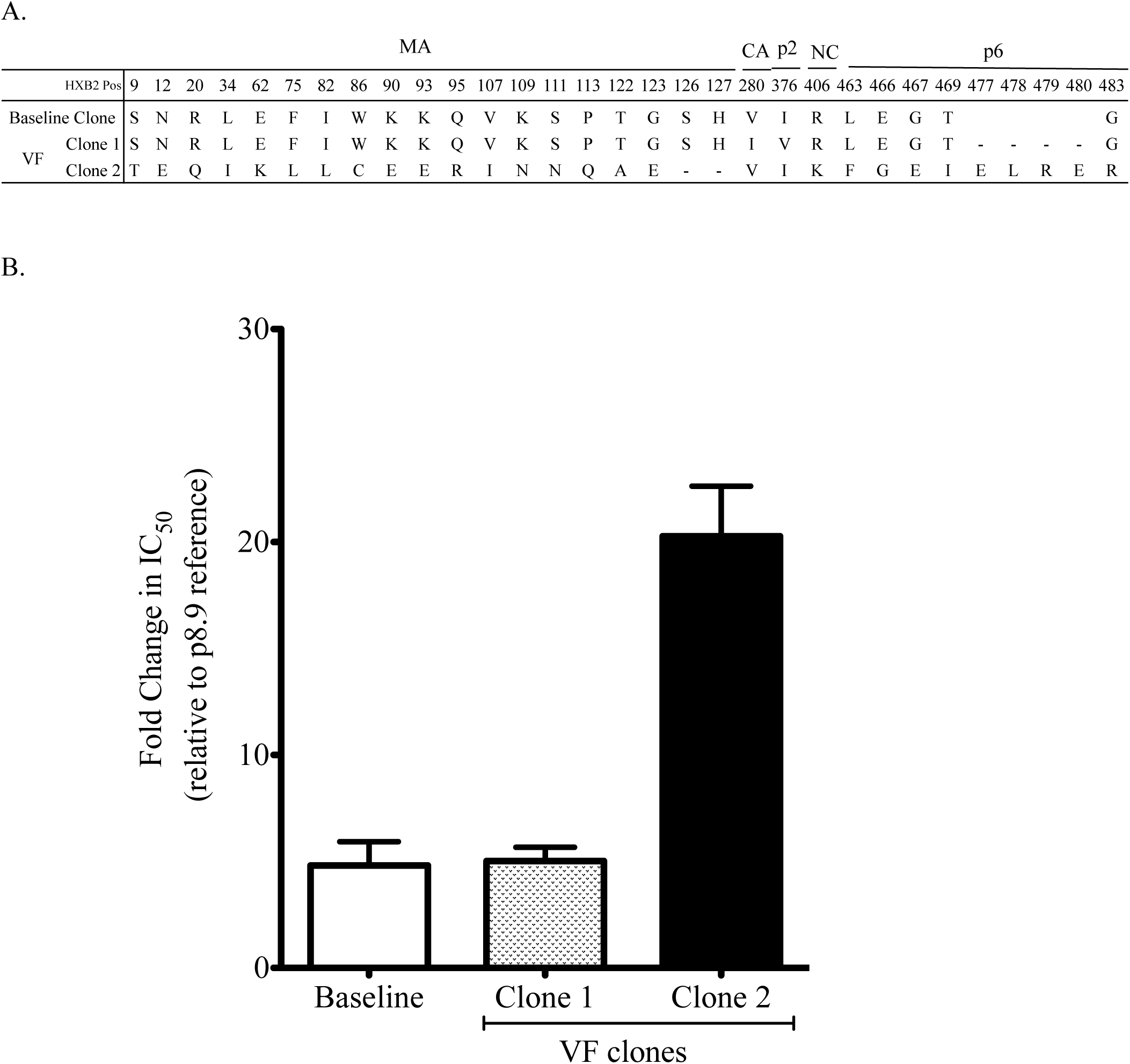
Variation in phenotypic PI susceptibility of full-length Gag-protease from HIV-1 infected patient at different time points. (A) is the sequences of the viral clones showing the amino acid changes in the MA, CA, P2, NC, p1 and p6 regions of gag between baseline (pre-PI treatment) and Viral failure (during PI treatment) timepoints. (B) Full-length Gag-protease was amplified from plasma samples and cloned into p8.9NSX+. VSV-g pseudotyped viruses encoding luciferase were produced by co-transfection in 293T cells. PI susceptibility of pseudovirions derived from each patient was determined using a single replication-cycle drug susceptibility assay as measured by luciferase activity. Data displayed are fold difference in IC50 values of LPV in comparison to that of the assay reference strain, p8.9NSX. Error bars represent the standard error of the mean of at least three independent experiments performed in duplicate.

We further explored this participant in order to elucidate determinants of resistance. Sequence alignment of full length gag and protease genes from sensitive and resistant clones revealed: 19 amino acid changes in matrix (MA), one change in each of capsid (CA), p2 and nucleocapsid (NC) as well as an insertion of four amino acids (E, L, R and E) in the p6 region of the resistant clone from *gag* position 477 (Figure 1). In protease, there was an M46V mutation in the resistant virus (Supplementary figure 2).

Interestingly, the VF sample was taken 64 months after PI initiation when the viral load was 66,277 copies/ml, and within this plasma sample two distinct virus clones were isolated (Figure 1, hatched and black bars). There was a 4-5 fold difference in LPV susceptibility between the two clones, suggesting a mixture of susceptible and ‘resistant’ viruses at the failure time point (Figure 1). We proceeded to map determinants of susceptibility using these two clones identified at failure. First, we sought to determine the role of a four amino acid insertion in the p6 domain. Using standard site directed mutagenesis techniques, amino acids E, L, R and E were inserted into a susceptible clone at position 477 in the p6 domain (Supplementary figure 3). Conversely, E, L, R and E residues were deleted in the less susceptible clone from the same location. There was no significant change in susceptibility to LPV as a result of the ELRE insertion (Supplementary figure 3).

### Matrix deletion of S126 and H127 confer reductions in LPV susceptibility

Given that the greatest number of changes occurred in the MA region, we sought to explore a possible role for MA amino acid changes on PI susceptibility. First, sequence changes occurring near the MA/CA cleavage site (within 10 amino acids) were considered. We noted that the more resistant virus had a deletion of Gag positions 126 and 127 as well as adjacent T122A and G123E mutations. Using site directed mutagenesis, serine (Gag position 126) and histidine (Gag position 127) residues were deleted in susceptible clone. Conversely, serine and histidine residues were inserted in the less susceptible clone. Deletion of *Ser* 126 and *His* 127 in the susceptible virus, led to a significant decrease in LPV susceptibility for the mutant virus (Figure 2). Conversely, the insertion of Ser and His residues in the resistant virus increased susceptibility of the mutant (Figure 2). However, the changes at positions 126 and 127 did not completely account for the differences in LPV susceptibility.

**Fig 2:**
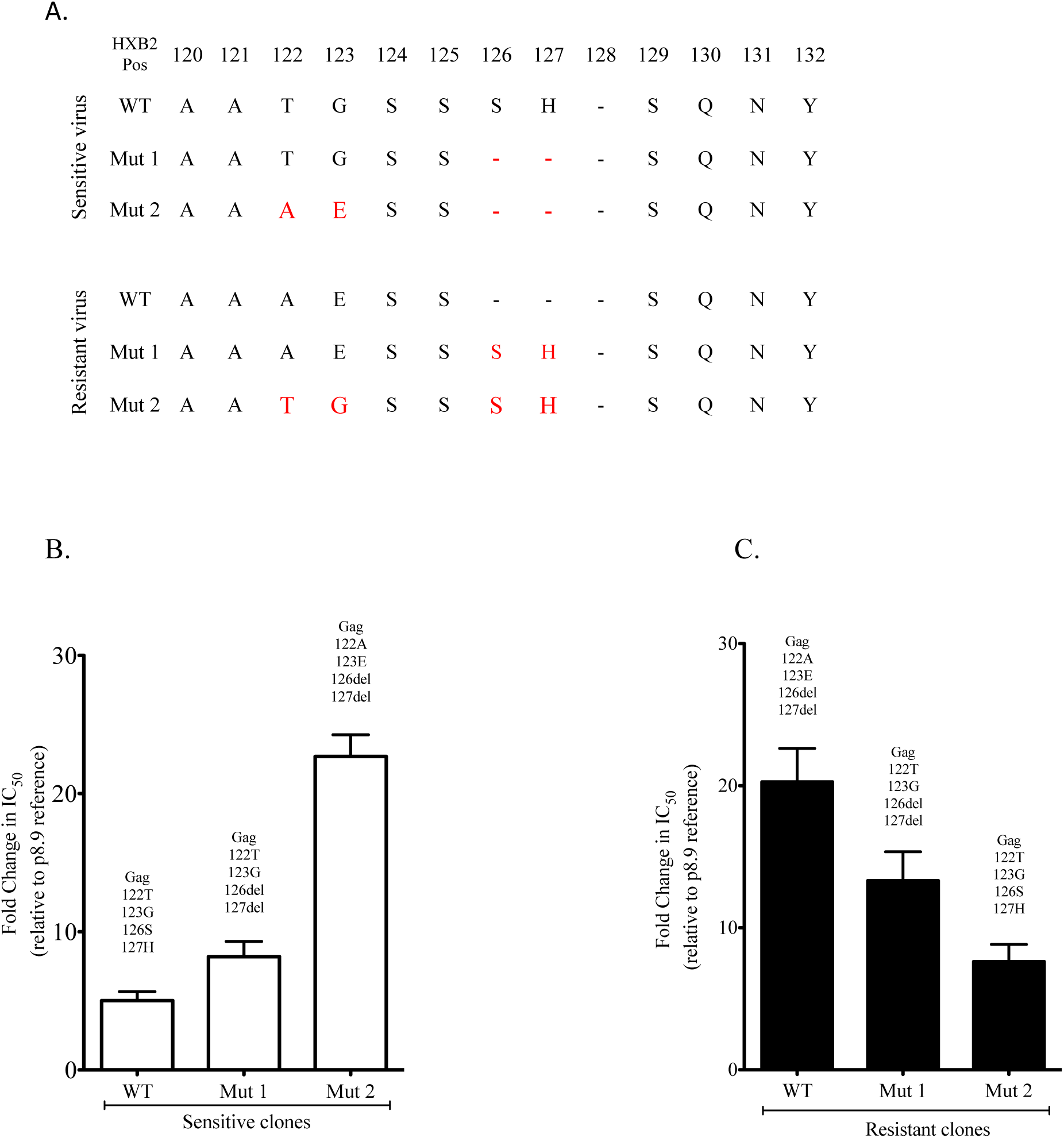
Gag 126del and 127del mutations occurring with T122A and G123E confers resistance to the protease inhibitor lopinivir in the absence of any major protease mutations. (A) sequences of the viral clones showing the amino acid changes (in red) introduced using standard site directed mutagenesis techniques. (B) and (C) Full-length Gag-protease with indicated mutations was amplified from plasma samples and cloned into p8.9NSX+. VSV-g pseudotyped viruses encoding luciferase were produced by co-transfection in 293T cells. PI susceptibility of pseudovirions derived from each patient was determined using a single replication-cycle drug susceptibility assay as measured by luciferase activity. Data displayed are fold difference in IC50 values of LPV in comparison to that of the assay reference strain, p8.9NSX. Error bars represent the standard error of the mean of at least three independent experiments performed in duplicate.

### The matrix deletions at S126 and H127 act synergistically with T122A and G123E gag

A combination of S126del, H127del and T122A, G123E mutations in the susceptible virus led to a 4x decrease in susceptibility to LPV (FC IC50 from 5.3 to 22.7), figure 2 and supplementary table 1. Conversely, S126*Ins*, H127*Ins* and A122T, E123G in the LPV resistant virus, led to a three-fold decrease in resistance as shown in Figure 2. We also tested the effect of the four amino acid signature on susceptibility to the second generation PI darunavir (DRV), and found no significant impact (Supplementary figure 4).

We sought to establish the effect of each of the four amino acid changes occurring alone. Using the resistant viral clone, four different mutant viruses were created with single amino acid changes at Gag: A122T, E123G, S126ins and H127ins. Results of the phenotypic drug susceptibility testing of these mutants showed that only E123G appeared to increase susceptibility (supplementary figure 5), and the combination of four amino acids had the greatest impact on LPV susceptibility.

We next tested whether the four amino acid signature T122A/ G123E/ S126del/ H127del could confer LPV resistance in a different subtype context. We chose the reference p8.9NSX subtype B virus and made the amino acid deletions at Gag positions 126 and 127 as well as the adjacent T122A and G123E mutations. In addition, we added a V128 deletion given that subtype CRF02_AG consensus contains this deletion as compared to subtype B. The five mutations (T122A/ G123E/ S126del/ H127del/V128del) reduced susceptibility to LPV by more than 3-fold, indicating that they are effective in a divergent subtype (Figure 3).

**Figure 3:**
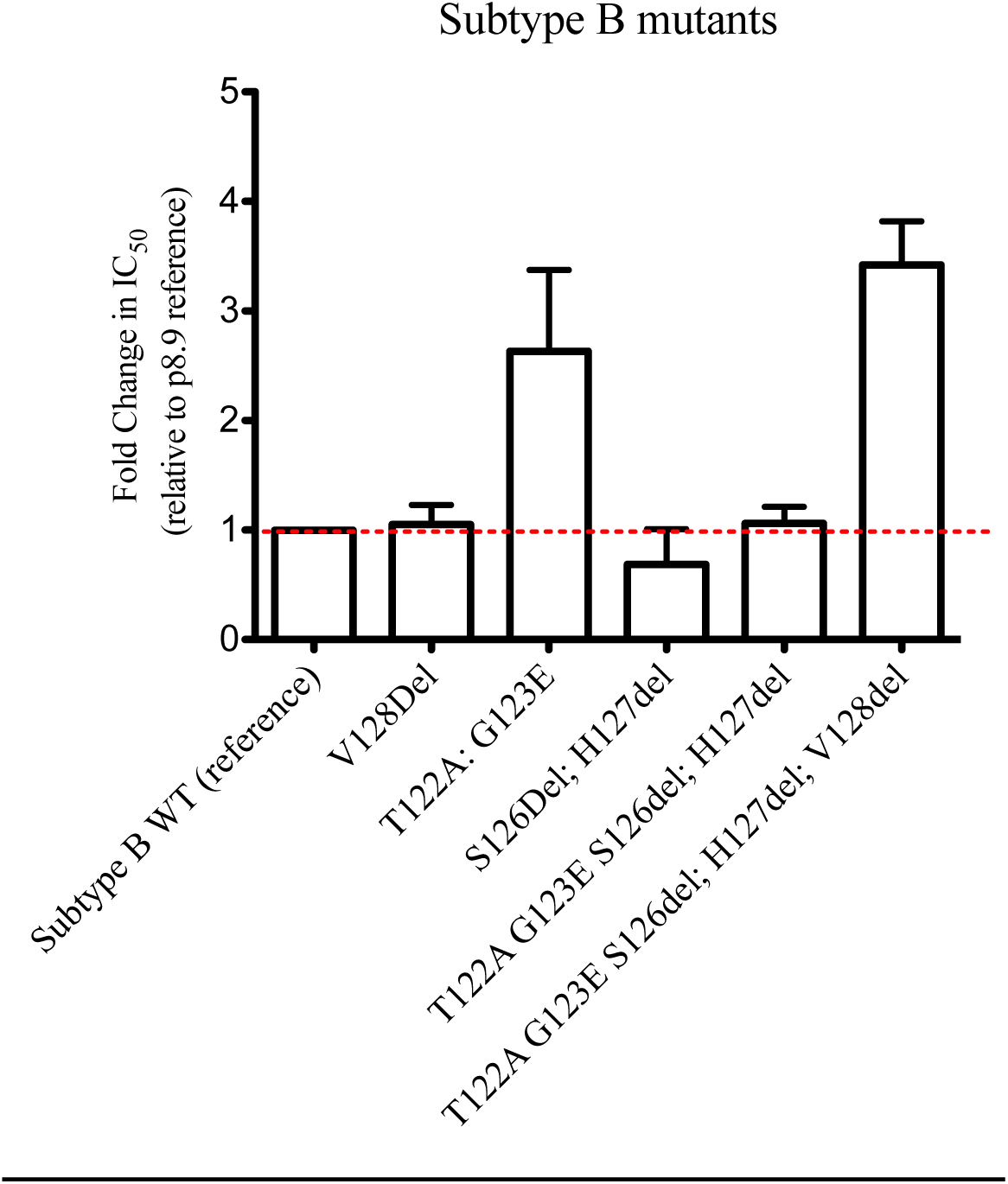
The four amino acid MA mutant signature can be introduced into subtype B to reduce PI susceptibility. Site directed mutants were generated in the subtype B reference strain used in our assays. The V128del was also added as this deletion is present in HIV-1 CRF02_AG. Data displayed are fold difference in IC50 values of LPV in comparison to that of the assay reference strain, p8.9NSX. Error bars represent the standard error of the mean of at least two independent experiments performed in duplicate.

### Matrix/Capsid (p17-p24) cleavage does not explain differential PI susceptibility

We hypothesized that the efficiency of MA/CA cleavage of HIV-1 polyproteins would differ between the susceptible and resistant clones in the presence of LPV. To test this hypothesis, we employed western blot analysis. Gag cleavage patterns were examined using the supernatants and cellular extracts of 293T cells transfected with each plasmid in the presence and absence of increasing concentrations of LPV (Figure 4). We probed with a polyclonal p24 antibody and as expected there was incomplete cleavage of p24-p2 at higher LPV doses in both virus containing supernatants and the cell extracts, consistent with previous data(30). We calculated ratios of p24/p41 to specifically probe the p17/p24 cleavage site in the vicinity of the four amino acid signature. The ratios did not indicate that resistant viruses were able to maintain cleavage at the p17/p24 site more efficiently than the sensitive virus in presence of LPV.

**Figure 4:**
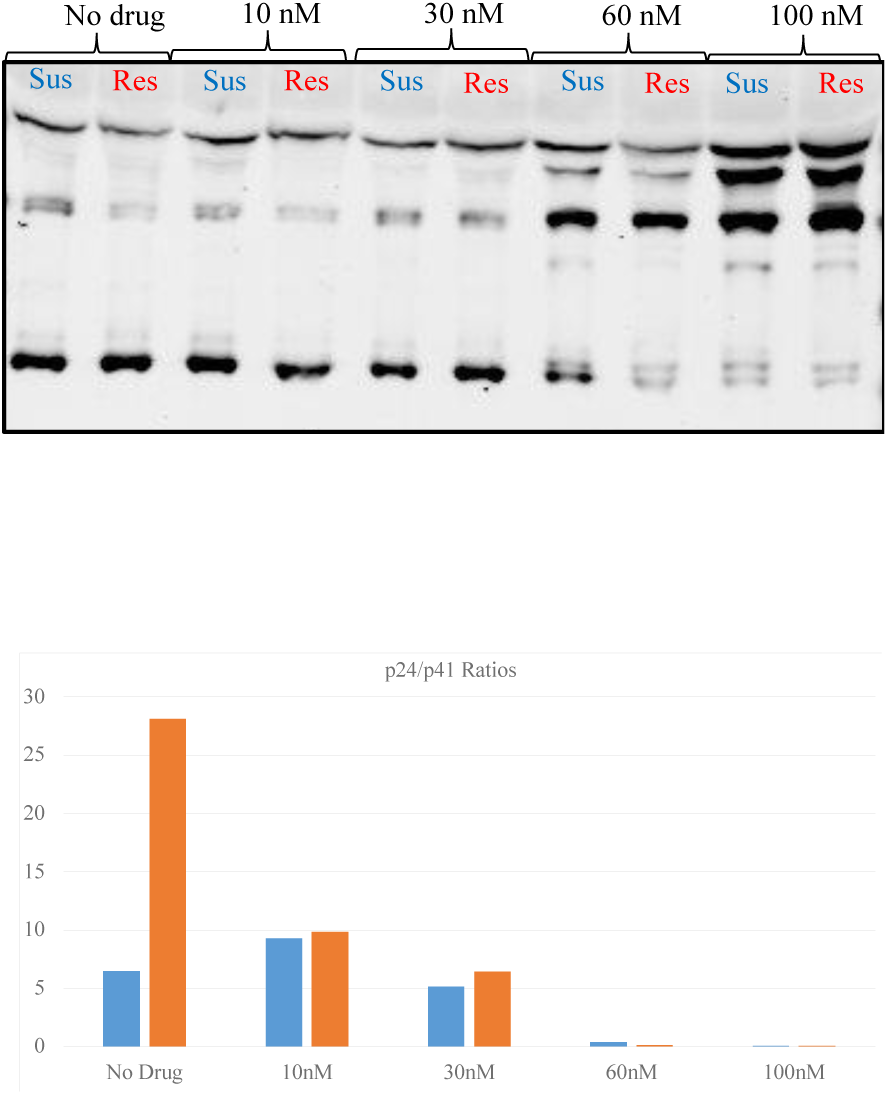
HIV-1 Gag cleavage efficiency in resistant versus susceptible isolates. Western blotting of virus containing supernatant, using a p24 antibody. In blue is the susceptible clone and orange the resistant clone. Ratios of p24/p41 are presented below and data are representative of 2 independent experiments.

### The resistance signature arises from a minority viral population detected at baseline

We proceeded to ask the question of when resistance emerged. Given the lengthy time period of over 5 years between the two samples we ideally needed a sample from an intermediate time point. We were able to identify a plasma sample on second line from 41 months, with VL of 241, 894. We refer to the 41-month time point as VF1 and the original 64-month time point as VF2. NGS analysis at whole genome level was undertaken for all 3 time points and table 2 shows variant frequencies at sites in Gag and Pol associated with drug exposure. Of note we observed loss of mutations to lamivudine (M184I), tenofovir (K65R) and efavirenz (K103N) between baseline and VF1. The individual was prescribed lamivudine, zidovudine and lopinavir/ritonavir for second line and the resistance data indicate lack of drug pressure from lamivudine.

**Table 2:** NGS variant derived data for three time points during LPV treatment showing the % of reads encoding resistance associated mutations in RT and Gag mutations known to be associated with protease inhibitor exposure from prior reports.

| **Gene** | **Mutation** | **Baseline**<br>**(0 months)**<br>**405,158 reads** | **VF1**<br>**(41 months)**<br>**250,932 reads** | **VF2**<br>**(64 months)**<br>**604,157 reads** |
| --- | --- | --- | --- | --- |
| Gag | E12K | 5% | 21% | 42% |
| Gag | R76K | 0 | 0 | 2% |
| Gag | Y79F | 0 | 0 | 2% |
| Gag | T122A | 4.8% | 13% | 26% |
| Gag | G123E | 5.0% | 13% | 26% |
| Gag | V128del | 100% | 100% | 100% |
| Gag | V370A | 5% | 0 | 1% |
| Gag | S373T | 97% | 98% | 100% |
| Gag | R409K | 3% | 0 | 1% |
| Gag | S451T | 100% | 1.7% | 0 |
| RT | K65R | 98% | 0 | 0 |
| RT | K103N | 94% | 0 | 0 |
| RT | E138K | 0 | 0 | 4% |
| RT | Y181C | 100% | 0 | 0 |
| RT | M184I | 100% | 1.2% | 21% |
| RT | M184V | 0 | 0 | 79% |

The NGS showed that T122A/ G123E were present at low abundance before initiation of PI (approx. 5% of reads, table 2). The proportion of T122A/ G123E increased at VF1 to 13%. These mutations were observed at increased frequency at VF2 both by target enriched NGS and also direct *gagpro* PCR from plasma, but NGS also showed emergence of lamivudine resistance mutant M184V, suggesting improved adherence to lamivudine between VF1 and VF2.

We next generated whole genome haplotypes for each time point using NGS data in order to firstly establish the phylogenetic relationships between viruses with differing PI resistance associated mutations, and also to determine the co-receptor usage of virus haplotypes as this might provide clues as to origins of virus variants (Figure 5). All inferred haplotypes were predicted to use CCR5 with FPR of <5%, and no CXCR4 using viruses were predicted in either of the two algorithms used.

**Figure 5:**
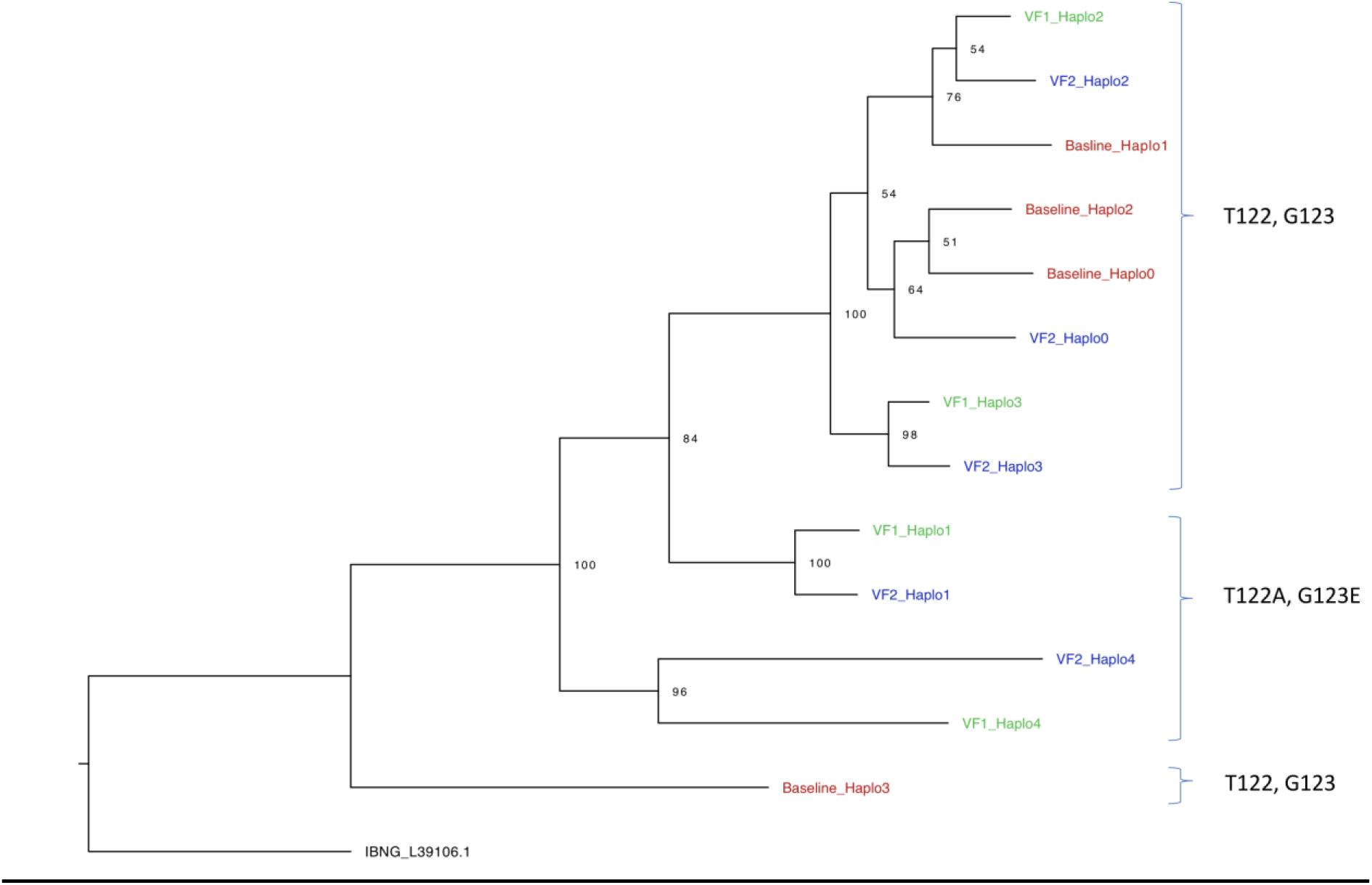
Whole genome HIV haplotype reconstruction using target enriched NGS. Illumina MiSeq data from each time point (baseline, VF1 and VF2), with maximum likelihood analysis and bootstrap support indicated using 1000 replicates. Labelled on the right side are the amino acids at positions Gag 122 and 123.

We proceeded to clone sequences from plasma at VF1 in addition to those previously cloned from VF2 and inferred phylogenetic trees. None of the four *gagpro* clones from baseline (before initiation of PI) contained any of the four amino acid changes T122A/ G123E/ S126del/ H127del, consistent with NGS data showing these variants were present at <5%(table 2). Clones from the intermediate time point VF1 clustered with the VF2 clones rather than with the baseline clones (Figure 6). Overall, there was excellent concordance between the inferred whole genome haplotypes and *gagpro* clones, though there appeared to be greater diversity in haplotypes. In vitro phenotypic drug susceptibility of cloned sequences revealed both sensitive and resistant viruses at VF1 as well as VF2 (Figure 6), with the resistant clones from VF1 and VF2 clustering together and sharing the 4 amino acid resistance associated signature S126del, H127del and T122A, G123E. As expected, the susceptible clones from VF1 and VF2 time points also clustered with each other in a distinct part of the tree.

**Figure 6:**
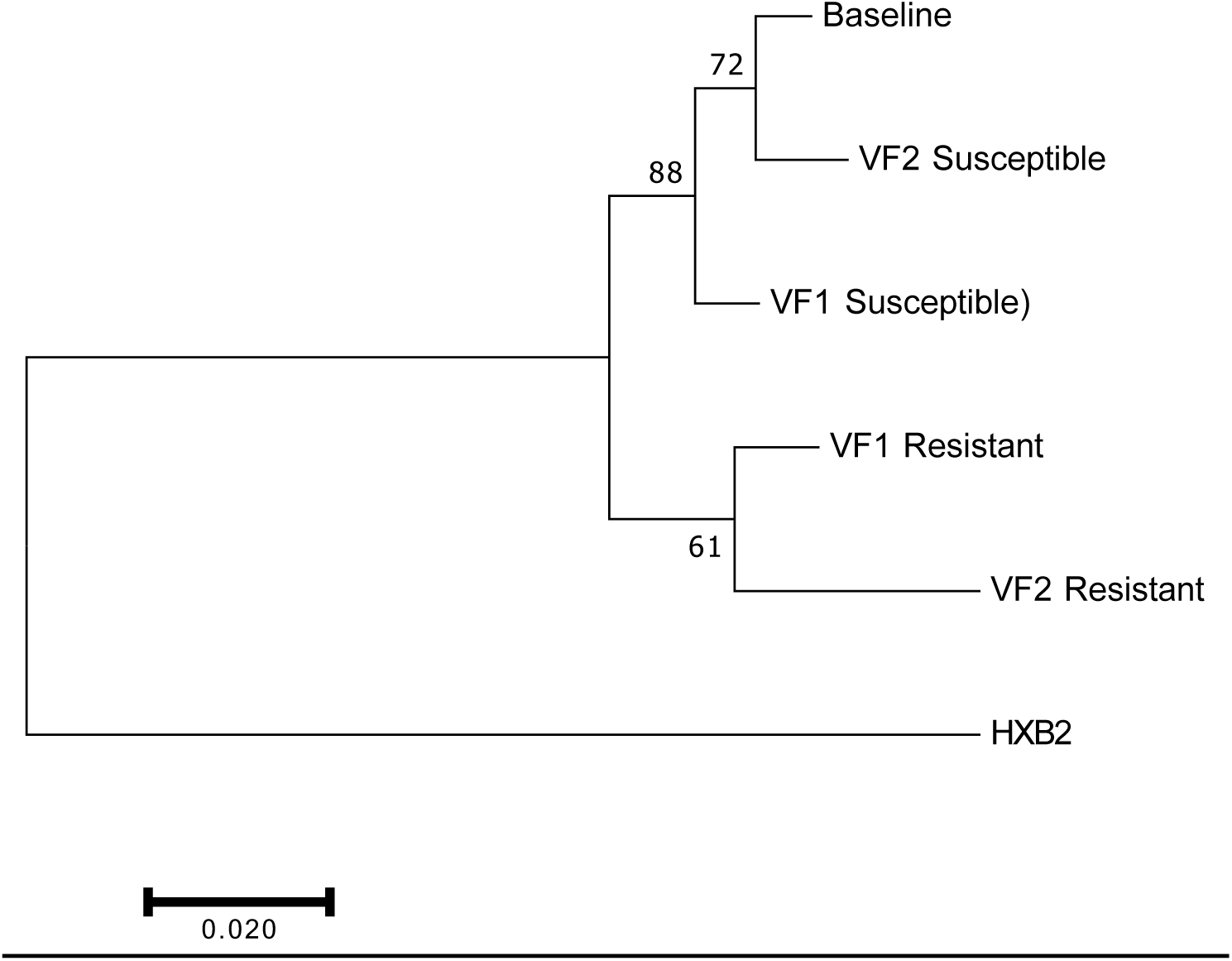
Phylogenetic relationships between viral Gag-protease plasma derived sequences isolated at baseline (pre-PI) and at two failure time points (VF1 and VF2). Illustrated is a maximum likelihood tree with bootstrap support indicated at nodes. Outlier indicated is HXB2, a subtype B virus. VF1: viral failure 1 time point 41 months post initiation of protease inhibitor therapy; VF1: viral failure 2 time point 64 months post initiation of protease inhibitor therapy.

### Persistence of both resistant and susceptible viruses can be explained by replication capacity

A surrogate for fitness in our assay is single round infectivity (measured in RLU) in the absence of drug, which is given a value of 100% for our reference subtype B virus. We measured the single round infectivity (replication capacity, RC) of clones bearing patient derived *gag-protease* sequences from each time point. Interestingly, resistant clones had a lower RC than susceptible viruses (around two fold), regardless of whether they were isolated from VF1 or VF2 (Figure 7). The mixture of sensitive and resistant strains is consistent with incomplete drug adherence and therefore variable drug pressure, or alternatively with compartmentalisation of virus sequences in anatomical areas with differing drug levels.

**Figure 7:**
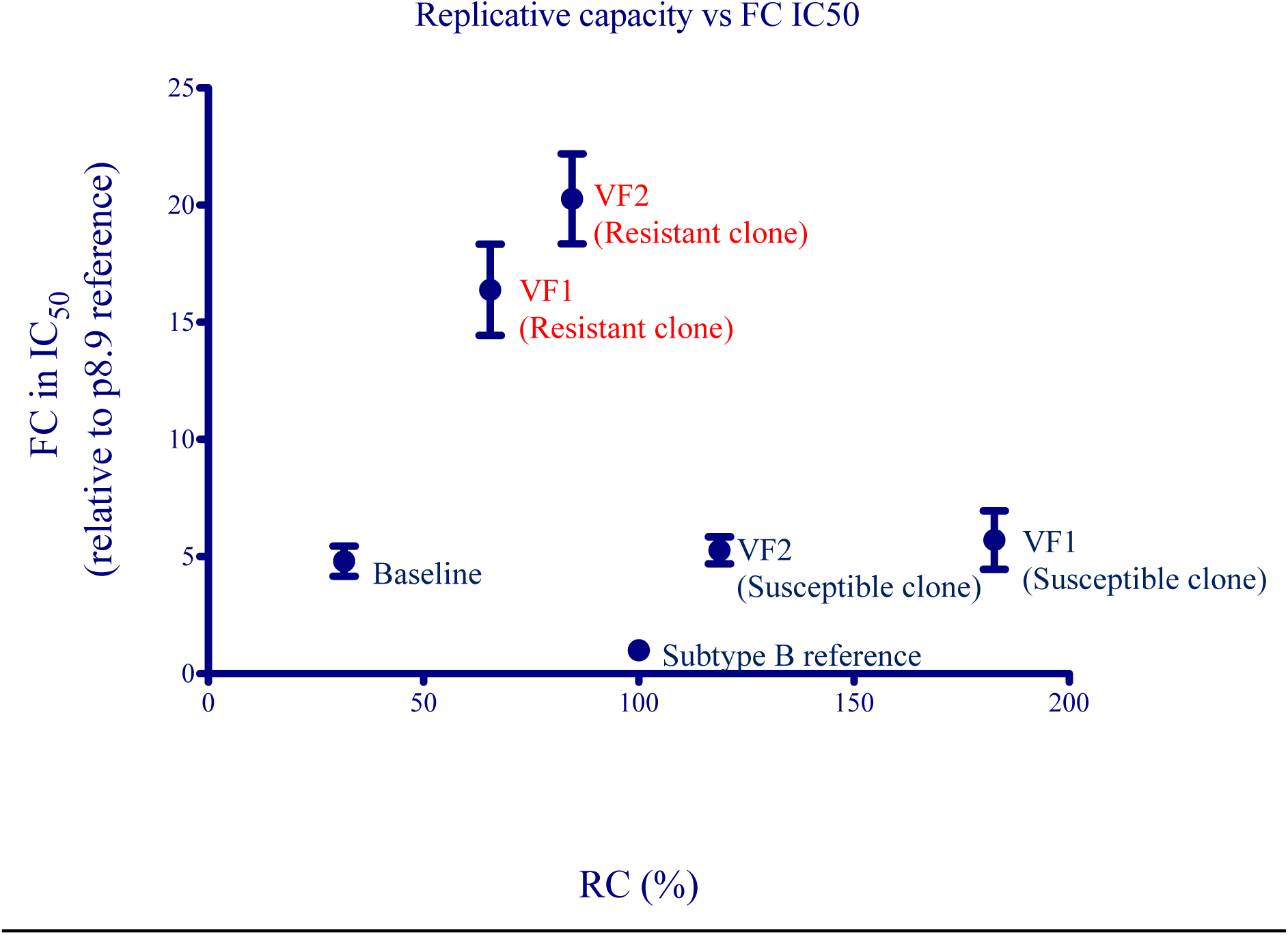
Relationship between single round infectivity (RC) and LPV susceptibility (FC in IC50) for patient derived viruses. Error bars represent the standard error of the mean of at least two independent experiments performed in duplicate.

## Discussion

Based on NMR and X ray crystallography studies, p17 comprises five major alpha helices connected primarily by short loops(33, 40). The C terminus of matrix is predicted to be disordered, which has hampered efforts to characterise the structural characteristics of this region. One study suggested that deletions at 125 and 126 would stabilise p17(41), indicating that despite disorder, changes in the region might lead to significant changes in stability and therefore possibly altered effects of protease inhibition on cleavage.

In this study on CRF02_AG and subtype G clinical isolates from a Nigerian cohort, we demonstrated the role of p17 amino acid mutations occurring near the p17/p24 cleavage site in contributing to PI resistance. The double deletion of *ser* and *his* at Gag 126 and 127 respectively had a modest impact on in-vitro phenotypic PI-susceptibility. When this deletion occurred together alongside T122A and G123E, we observed a 4-5 fold decrease in susceptibility to lopinavir. The four mutation combination was also able to confer similar resistance to a subtype B virus, indicating that it may emerge across subtypes.

G123E was reported to arise when viruses were propagated with investigational protease inhibitors KNI-272 and UIC-94003(42). Gag G123E was found to potentially interact with protease by NMR (33), providing a potential mechanism for its effect. Our present study also implicates G123E in reduced PI susceptibility, but we show here that the combination of mutations that was observed in the patient was needed for maximal effect.

We aimed to understand the mechanism at play in the T122A, G123E, S126del, H127del phenotype. Western blotting of virus containing supernatants from producer cells, or the cells themselves, did not reveal significant differences in cleavage with or without drug at the p17-p24 cleavage site. Gag cleavage patterns at the MA/CA cleavage site (as obtained from p24/p41 ratios) were similar between the susceptible and resistant viral clones. Therefore, the rescue of infectivity in the presence of drug is not explained by a change in the detective cleavage pattern of Gag caused by inhibitor. This suggests that resistance is not mediated by rescuing wild type Gag cleavage patterns but rather by tolerating the changes that the inhibitor drives. This in turn suggests that the defect in cleavage may be indirectly related to the mechanism of inhibition and inhibition of infection may be mediated by more subtle effects than simply defects in overall cleavage levels. This more complex mechanism may be particularly important at drug concentrations at which defects in Gag cleavage measured by western blot are not apparent.

We used NGS to explore the dynamics of emergence of Gag amino acid changes during ongoing viremia under PI treatment. We were able to detect both T122A and G123 at low abundance at baseline, prior to PI exposure. Importantly, PCR from plasma RNA using *gagpro* specific primers did not amplify any sequences with these changes at baseline, highlighting an important contribution of NGS to the study of drug resistance. Whole genome reconstruction enabled us to infer phylogenetic trees and confirm findings that resistance conferring mutations occurred at both time points in phylogenetically related sequences. All virus haplotypes were predicted to be CCR5 using and therefore sensitive to the CCR5 antagonist maraviroc.

In future work it would be interesting and important to know whether Gag mutations are capable of facilitating emergence of major protease mutations in prolonged culture conditions under suboptimal drug pressure. This could potentially explain why prevalence of major protease mutations increases over time during PI exposure in clinical studies(43). Next one could perform population dynamics simulations to incorporate RC and susceptibility data in order to model the proportion of resistant and susceptible viruses over time, and possibly therefore predict emergence of major mutations in *protease*.

Our data are limited by the small sample size, lack of availability of plasma drug level measurements, and by the use of standard clonal approaches as opposed to single genome sequencing and amplification. Nonetheless, we hypothesise that the four amino acid HIV-1 Gag signature is a contributory factor in PI failure in this PLWH from Nigeria.

As we move towards next generation sequencing, this work highlights the limitations of current genotyping methods to infer PI susceptibility, and supports sequencing outside protease in to broaden the evidence base for the clinical management of patients who experience VF on PIs without major protease mutations. The work may ultimately also help to identify define individuals with lower PI susceptibility before treatment with this class of drugs.

## Transparency declarations

RKG has received speaker fees for ad hoc consulting from Gilead and ViiV. NN has received an investigator award grant from Gilead.

